# Functional screening of ZIP8 naturally occurring variants identifies pathogenic mutations and trafficking defects

**DOI:** 10.64898/2026.05.11.724455

**Authors:** Michael Nikolovski, Tianqi Wang, Aaron Sue, Keith MacRenaris, Hongyan Zhao, Thomas V. O’Halloran, Jian Hu

## Abstract

The rapid expansion of human genomic data has revealed a large number of naturally occurring variants, creating a major challenge for functional annotation. The human metal transporter SLC39A8 (ZIP8) is a clinically important, promiscuous divalent metal transporter, yet most of its documented variants remain uncharacterized. Here, we developed a workflow to functionally evaluate ZIP8 variants by integrating laser ablation inductively coupled plasma time-of-flight mass spectrometry (LA-ICP-TOF-MS) with scaled-up cell-based transport assays. Using this method, we systematically analyzed 33 naturally occurring missense variants located in the extracellular domain (ECD) of ZIP8. The assay enables direct quantification of intracellular metal accumulation with substantially improved throughput (∼150 samples per hour). Functional screening identified 14 potential pathogenic variants with significantly reduced transport activity. Comparison with computational predictions revealed a moderate correlation between activity and AlphaMissense pathogenicity scores (R^2^ = 0.423), while an error rate of ∼20% underscores the need for experimental validation. Flow cytometry analysis showed that most loss-of-function variants exhibit impaired trafficking of the protein to the cell surface possibly due to mutation-caused protein misfolding or instability. Structural mapping of activity-compromised variants, together with functional assessment of the ZIP8-ECD, highlights the importance of this domain in ZIP8 expression and intracellular trafficking. Together, this work establishes a scalable approach for functional screening of metal transporter variants and provides new insights into the structure–function relationships of ZIP8.

## Introduction

With the rapid progress in genome sequencing, a vast number of naturally occurring genetic variants have been identified across the human genome (1,2). While this wealth of genetic information provides unprecedented opportunities to understand gene function and disease mechanisms, it also presents a significant challenge regarding the functional annotation of the large number of uncharacterized variants (3). One prominent example is SLC39A8 (or ZIP8), which belongs to the LIV-1 subfamily of the Zrt/Irt-like protein (ZIP, or SLC39A) metal transporters (4-10). In humans, fourteen ZIP transporters (ZIP1–ZIP14) have been identified with nine of them in the LIV-1 subfamily. Compared with other human ZIPs, SLC39A8 (ZIP8) and its close homolog SLC39A14 (ZIP14) are more promiscuous, as they transport multiple divalent metals, including Zn^2+^, Mn^2+^, Fe^2+^, Co^2+^, Cd^2+^, Pb^2+^, and Cu^2+^ (11-15). Two clinical studies reported in 2015 first revealed that patients suffering from various neurological disorders exhibited severe Mn deficiency caused by loss-of-function mutations in *slc39a8* (16,17), thereby establishing a new genetic disorder, SLC39A8-congenital disorder of glycosylation (SLC39A8-CDG). Subsequent studies have identified and characterized additional pathogenic variants associated with various diseases (18-26), in particular those in the neurological system. Despite these advances, the number of experimentally characterized variants remains much smaller than the total number of SLC39A8 variations identified in the human population. To date, the UniProt database lists 369 missense variants but only eight missense variants have been experimentally confirmed as pathogenic based on biochemical evidence. Most documented SLC39A8 variations remain either unannotated or classified as variants of unknown significance (VUS). This situation highlights a substantial gap between the clinical need to understand the functional consequences of these variants and the limited availability of reliable functional annotations. Similarly, efforts to clarify the functional consequences of many naturally occurring mutations in other ZIPs are needed to uncover potential pathogenicity (27,28).

Computational approaches have been developed to help address this challenge. Tools based on sequence conservation, structural modelling, or machine learning, such as the recently developed AlphaMissense (AM) (29), aim to predict the pathogenicity of genetic variants. However, these approaches often show limited predictive accuracy and can yield inconsistent results across different methods. As shown in **Figure S1**, among the eight known pathogenic SLC39A8 variants, AM correctly predicts only four as pathogenic, while annotating one as ambiguous and three as benign. The Combined Annotation Dependent Depletion (CADD), another widely used metric for predicting functional impact of genetic variations (30,31), appears to perform better in predicting pathogenic variants when a loose cutoff is applied (**Figure S1**), but this is primarily due to the tendency to classify variants more frequently as pathogenic or ambiguous. The incorrect predictions for known pathogenic variants and the inconsistent predictions for many VUSs indicate current computational tools have room for improvement. . Experimental determination of variant function is crucial not only for correctly identifying pathogenic mutations but also for improving predictive algorithms through the availability of high-quality training data.

In this study, we present an experimental workflow to enable functional screening of ZIP8 variants by combining laser ablation inductively coupled plasma time-of-flight mass spectrometry (LA-ICP-TOF-MS) with a cell-based metal transport assay. This approach allows quantification of intracellular metal accumulation with substantially increased throughput. Using this method, we screened 33 naturally occurring variants located in the extracellular domain (ECD) of ZIP8 and identified 14 potential pathogenic variants. By integrating data from the activity measurements with flow cytometry analysis to quantify the cell surface and total expression levels, we determined that the reduced cation transport activity is primarily caused by inefficient trafficking of the protein to the cell surface for most variants. The screening results reveal sites that have structure-function implications for ZIP8-ECD.

## Results and Discussion

### Selection of ZIP8 naturally occurring variants

To evaluate the function of naturally occurring ZIP8 variants, we selected mutations based on their AM pathogenicity scores, with scores of 0-0.33 classified as benign, 0.34-0.56 as ambiguous, and 0.57-1.00 as pathogenic. In this study, we focused on variants located in the N-terminal extracellular domain (ECD) of ZIP8 (residue 25-124). Compared with the transmembrane domain, ECD is less conserved (**Figure S2**), which may lead to reduced accuracy for predictions by AM. It is because AM relies on evolutionary conservation as a key source of information and thus its predictive confidence and performance are often lower in less conserved regions (32). Based on these criteria, we selected a total of 33 of the 108naturally occurring missense variants in the ZIP8-ECD for functional assay (**Supporting data S1**), including 13 variants predicted by AM to be pathogenic, 4 predicted to be ambiguous, and 15 predicted to be benign according to the pathogenicity scores, together with one confirmed pathogenic variant (C113S) used as a positive control. The V33M mutation was initially reported to be pathogenic as it was identified in a compound heterozygous patient (17), but later biochemical studies showed conflicting results from no activity to increased activity (20,33-35). We therefore included V33M in this study to clarify whether V33M is a deleterious mutation.

### A work flow to measure transport activity of ZIP8 variants with enhanced throughput

Given the large number of variants to be functionally evaluated, we adopted and optimized a previously reported metal quantification method using LA-ICP-TOF-MS to analyse an array of cell samples (36) (**Figure 1A**). In brief, HEK293T cells were cultured in 96-well plates and transfected with an empty vector or vectors encoding wild-type ZIP8 or its variants. After 24 hours, cells were subjected to a metal transport assay using Cd^2+^ as the substrate. Cd^2+^ was selected because of its low natural background in cells. After the transport assay, cells were suspended and a small volume (∼7.5 μl) was spotted onto a siliconized glass slide to generate a sample array. With careful handling, a single slide can accommodate up to 50 samples. The array was air-dried and subsequently analyzed by LA-ICP-TOF-MS. Each droplet on the sample array was scanned by five parallel lines with a linewidth of 40 µM using laser and the evaporated gas was analyzed by ICP-TOF-MS to quantify ^111^Cd (one of the naturally occurring stable isotopes of cadmium) and ^31^P. The count ratios of ^111^Cd/^31^P are used to represent metal contents, as the variation in cell number has been levelled out using the phosphorus content (36).

**Figure 1.**
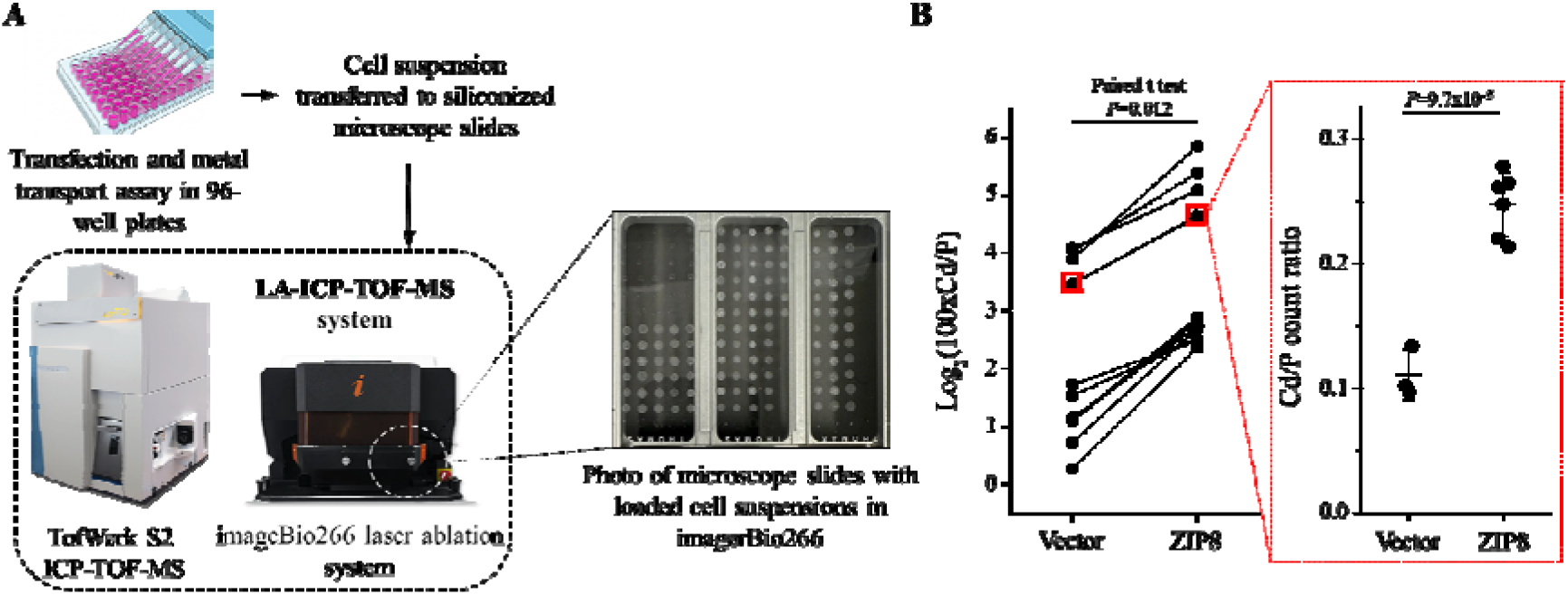
Measurement of ZIP8 transport activity with enhanced throughput. (**A**) Workflow of transport assay and metal quantification by LA-ICP-TOF-MS. (**B**) Method validation. *Left*: Statistical analysis for the vector group and WT ZIP8 group in 11 independent experiments. For clarity, the raw ^111^Cd/^31^P ratios are multiplied by 100 and then transformed using log2. Solid lines connect paired control and ZIP8 samples in each experiment. Two-tailed paired *t* test was used to examine the null hypothesis. *Right*: Raw ^111^Cd/^31^P ratios of the vector group and WT ZIP8 group from one representative experiment indicated with red boxes. Two-tailed *t* test was used to examine the null hypothesis.

This method was originally developed to study ZIP4 using cells grown in 24-well plates, we first tested whether it could be adapted for use with 96-well plates to study ZIP8. Across 11 independent experiments conducted on different days, we consistently observed a statistically significant increase in Cd accumulation (expressed as the count ratio of Cd/P) in cells expressing FLAG-tagged ZIP8 compared with cells transfected with the empty vector (**Figure 1B**, left). In each experiment, which included at least three biological replicates, Cd accumulation in the ZIP8 group was significantly higher than in the vector control group (**Figure 1B**, right). We therefore defined the difference between these two groups as ZIP8 transport activity. These results demonstrate that the LA-ICP-TOF-MS–based assay remains robust when cells are cultured and processed in 96-well plates. A major advantage of this approach is its substantially increased throughput. Compared with conventional ICP-MS analysis of liquid samples, the array-based LA-ICP-TOF-MS method can measure up to ∼150 samples per hour, representing up to a six-fold improvement in throughput. When combined with cell handling in 96-well plates using a multichannel pipette, the overall efficiency of the workflow is greatly enhanced.

### Functional screening ZIP8 variants and comparison with predictions

Independent batches of metal transport assays for all variants were conducted following the workflow established above. For each variant, at least three biological replicates were included in each experiment, and three independent experiments were performed. The results for all 33 variants are summarized in **Figure 2A**, where the transport activities of the variants are expressed as percentages relative to wild-type ZIP8. Among these variants, 17 exhibited significantly reduced activity, of which 15 showed activities below 70% of wild-type levels. These include the positive control C113S, which displayed 23 ± 7% of the wild-type ZIP8 activity. We classified variants with activity below 70% as pathogenic. This threshold was chosen based on the well-characterized pathogenic variant A391T, which has been reported to exhibit an approximately 30% reduction in activity compared with wild-type ZIP8 (26). V33M, a mutation identified in a compound heterozygous individual (17), resulted in an activity comparable to wild-type ZIP8 (99%), supporting its classification as benign (34). In contrast, S44W, another mutation also identified in a compound heterozygous individual (22), showed markedly reduced activity (34%), indicating a potential pathogenic mutation (33). N72 is a known glycosylation site of ZIP8 (37), and the unchanged activity of the N72S variant indicates that *N*-glycosylation at this position is consistent with the notion that *N*-glycosylation is dispensable for ZIP8-mediated metal transport (38).

**Figure 2.**
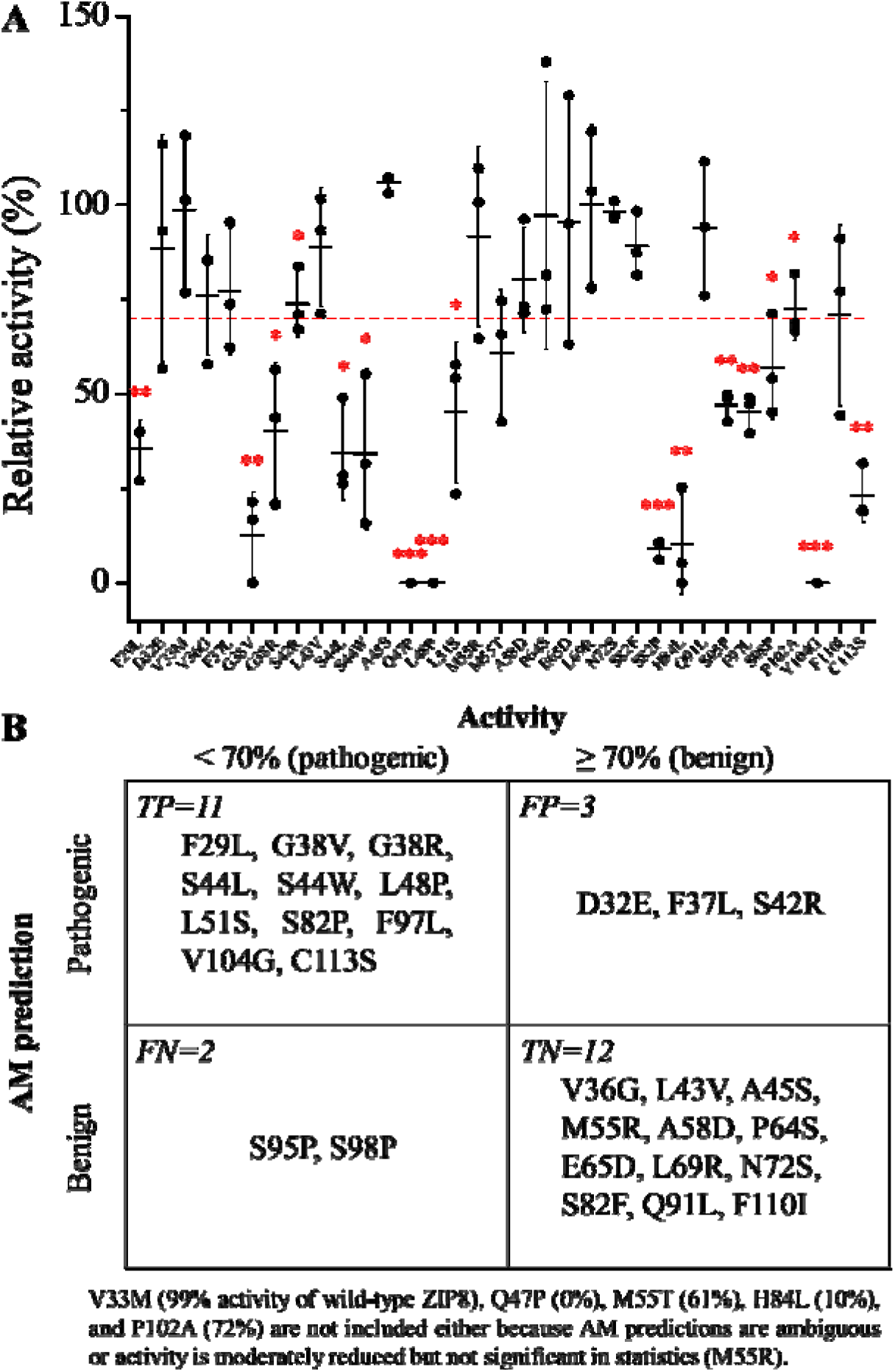
Transport activities of 33 variants in ZIP8-ECD. (**A**) Transport activities of variants expressed as percentages relative to wild-type ZIP8. The three data points for each variant are the mean values obtained from three independent experiments. In each independent experiment, at least three biological replicates were used to calculate the mean relative activity of each variant. The horizontal lines and error bars indicate the mean and standard deviation (S.D.), respectively. The red dashed line indicates 70% relative activity, which is used in this study as the threshold to classify variants as pathogenic or benign. Two-tailed t test was used to examine null hypothesis. ^*^: *P*<0.05; ^**^: *P*<0.01; ^***^: *P*<0.001. (**B**) Confusion matrix comparing AM predictions with classifications experimentally determined in this study. TP, true positive (correctly identified pathogenic variants); TN, true negative (correctly identified benign variants); FP, false positive (incorrectly identified pathogenic variants); FN, false negative (incorrectly identified benign variants).

Based on these results, we compared the experimentally determined transport activities with the AM pathogenicity scores for each variant. A moderate negative linear correlation was observed (*R*^2^ = 0.423, *P* = 4.2 × 10^-5^; **Figure S3**), indicating that variants with higher predicted pathogenicity tend to exhibit lower transport activity. In comparison, CADD scores also showed a negative correlation with transport activity, but with weaker predictive power (*R*^2^ = 0.240, *P* = 0.0039). These results suggest that AM outperforms CADD in predicting the functional impact of ZIP8 variants in this dataset. To further evaluate the performance of AM in classifying pathogenic variants, we constructed a confusion matrix (**Figure 2B**). Variants with ambiguous AM scores (0.34-0.56) were excluded from this analysis. Based on this classification, the accuracy, specificity, sensitivity, and false discovery rate of AM prediction were calculated to be 82%, 80%, 79%, and 21%, respectively. These performance metrics are comparable to those reported in studies involving larger datasets (39,40), supporting the reliability of AM for predicting variant function, while also highlighting the importance of experimental validation, given an error rate of approximately 20% in this study.

### Examining trafficking efficiency of potentially pathogenic variants

Pathogenic mutations often cause protein dysfunction by disrupting protein folding and stability. For membrane transporters, misfolding or structural instability frequently leads to reduced protein expression and/or defects in trafficking from the endoplasmic reticulum (ER) to the plasma membrane. To examine whether any of the identified potential pathogenic variants exhibit such defects, we employed flow cytometry to compare both the total expression and cell surface expression of FLAG-tagged variants in HEK293T cells with those of wild-type ZIP8 (**Figures 3A & S4**). With a few exceptions, most variants exhibited moderately reduced total expression levels compared with wild-type ZIP8. The reduction in cell surface expression was generally more pronounced, as indicated by the observation that all variant data points fall to the right of the line connecting the wild-type ZIP8 data point and the origin of the plot in **Figure 3A**. When trafficking efficiency, which is defined as the ratio of relative surface expression to relative total expression (39,41), was calculated and compared with that of wild-type ZIP8, all variants except two showed significantly reduced trafficking efficiency (**Figure 3B**). Notably, five variants (G38V, Q47P, L48P, S82P, and V104G) exhibited particularly low trafficking efficiency, suggesting severe defects in protein folding and/or intracellular trafficking. In contrast, the remaining variants displayed only moderate reductions in trafficking efficiency.

**Figure 3.**
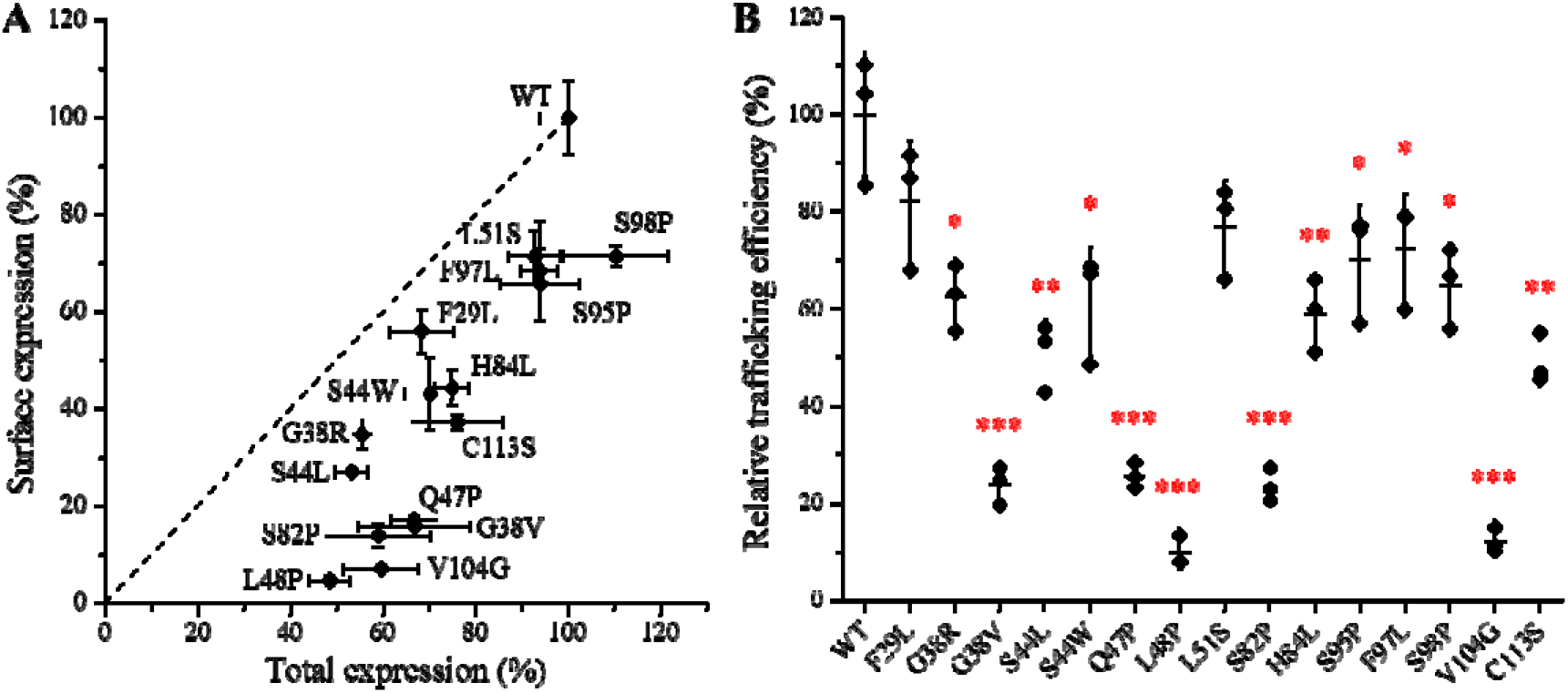
Total and cell surface expression of FLAG-tagged ZIP8 variants determined by flow cytometry. (**A**) Plot of relative cell surface expression against relative total expression for ZIP8 variants with activities below 70% of wild-type ZIP8. The data shown are from one of two independent experiments, with error bars represensting one S.D. of three biological replicates. Wild-type ZIP8 and the variants are labeled. The corresponding raw data are shown **Figure S4C.** The dashed line connects the data point for wild-type ZIP8 and the origin of the plot. (**B**) Trafficking efficiencies of the variants relative to wild-type ZIP8. Trafficking efficiency is defined as the ratio of relative surface expression to relative total expression shown in (A).

### Classification of potential pathogenic variants

Next, we sought to explain the reduced transport activities observed for the identified potential pathogenic variants. To this end, we examined the relationship between transport activity and trafficking efficiency for each variant. This analysis allowed us to classify the variants into three groups (**Figure 4**). The largest group comprises eight variants that exhibit moderately reduced transport activity (35–60% of wild-type ZIP8) together with moderately reduced trafficking efficiency (50–80% of wild-type levels). The second largest group includes the five variants that display very low cell surface expression and severely impaired trafficking efficiency. The positive control, C113S, falls between these two groups but is closer to the first group. In contrast, H84L appears to be an outlier. Although this variant shows only moderately reduced trafficking efficiency (∼60% of wild-type ZIP8), its transport activity is drastically reduced (∼10% of wild-type levels). Overall, these results suggest that for most variants with reduced transport activity, impaired trafficking, which is likely resulting from defects in protein folding or stability, accounts for the majority of the functional loss. In contrast, the severe activity reduction observed for H84L cannot be fully explained by its moderate trafficking defect, indicating that this mutation likely impairs transport efficiency through a distinct and yet unknown mechanism.

**Figure 4.**
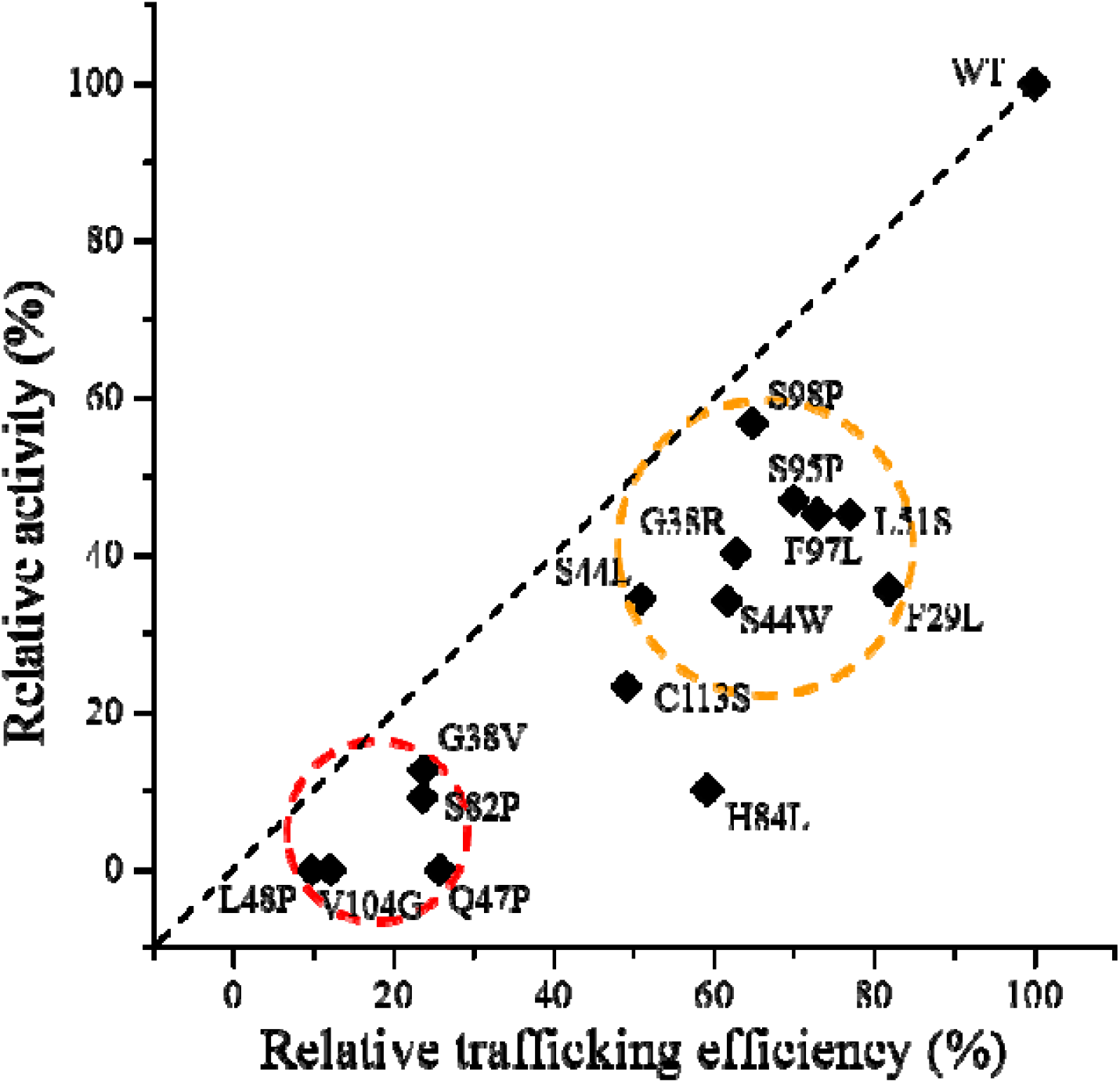
Classification of potential pathogenic variants. Relative activity of each variant is plotted against relative trafficking efficiency, revealing two major groups with H84L as an outlier. The larger group (in the orange circle) comprises variants with moderate reductions in both activity and trafficking efficiency, whereas the smaller group (in the red circle) includes variants with severely reduced activity and trafficking efficiency. The positive control C113S is near the larger group. In contrast, the H84L variant exhibits only a modest reduction in trafficking efficiency but a severe reduction in activity and does not belong to either group.

### Implications on structure-function relationship of ZIP8-ECD

Functional screening of naturally occurring mutations inZIP8-ECD provides important insights into its structure-function relationships, as revealed by mapping these mutations onto the structural model of ZIP8 dimer (**Figure 5**). A hallmark of the LIV-1 subfamily is ECD dimerization with a conserved PAL-like motif sitting at the center of the dimerization interface (42). In ZIP8, this motif corresponds to residues ^102^PAV^104^ (Figure 6B), where two variants, P102A and V104G, were examined in this study. The P102A variant exhibited a significant but modest reduction in activity (72% of wild-type ZIP8; **Figure 2**). In contrast, the V104G variant showed no detectable transport activity and severely impaired trafficking efficiency, despite only a moderate reduction in total expression. This observation is consistent with findings for ZIP4, where replacement of the PAL motif with an AAA sequence results in complete loss of activity and an approximately 80% reduction in cell surface expression. Another variant with severely diminished activity, H84L, is also located at the dimerization interface. The polar side chain of H84 is surrounded by hydrophobic residues from both protomers (**Figure 5**, upper right panel), and how substitution with leucine nearly abolishes activity remains unclear and warrants further structural investigation. Together, these results underscore the importance of ZIP8-ECD dimerization in maintaining transporter function.

**Figure 5.**
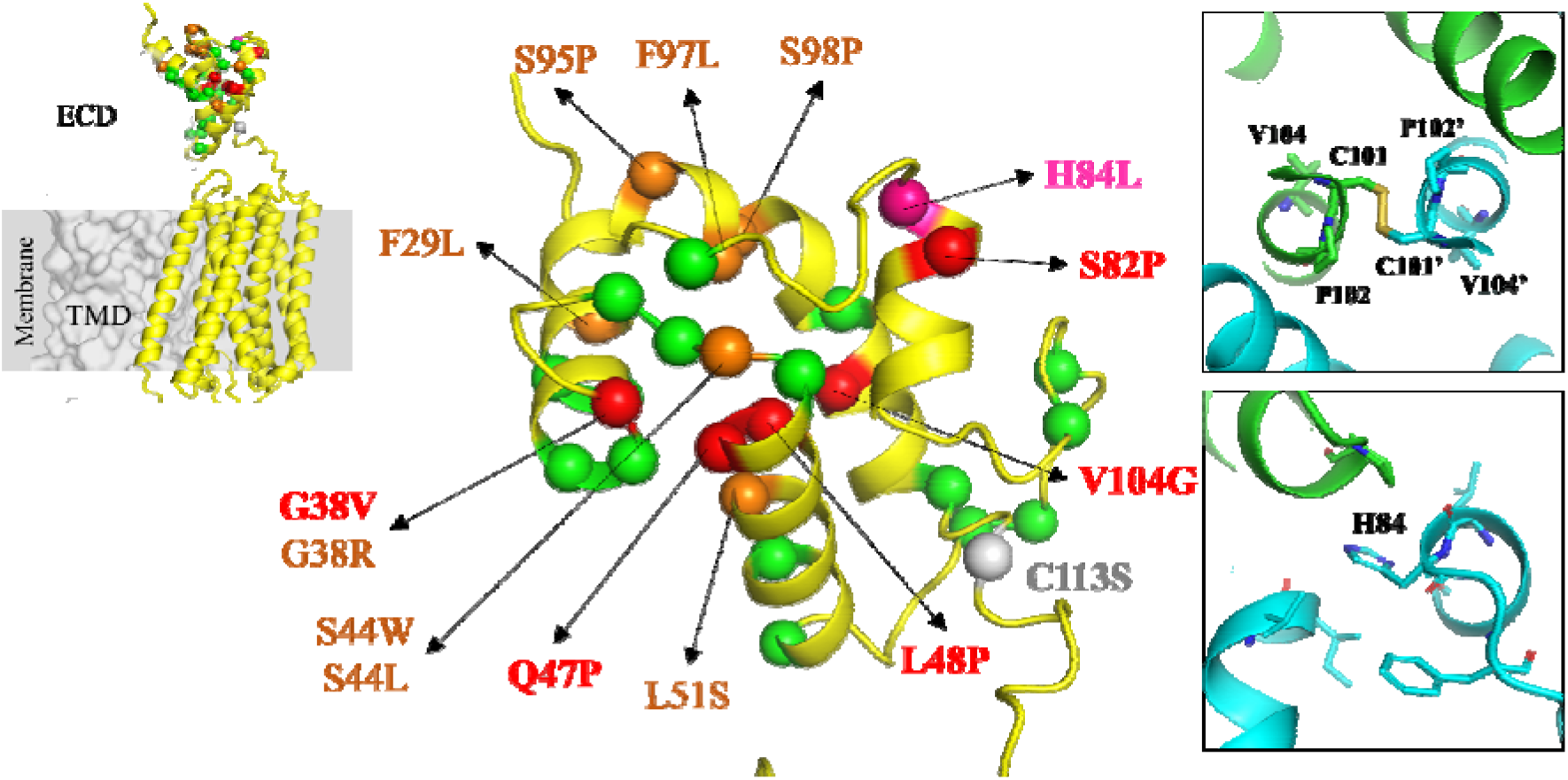
Mapping naturally occurring variants studied in this work onto a structural model of human ZIP8. The ZIP8 dimer structure (upper left inset) was generated by AlphaFold 3, with one monomer shown in cartoon mode (yellow) and the other in surface mode (grey). The ECD dimer is shown in the center. The Cα atoms of variant residues are shown as spheres and colored as follows: green, benign; orange, potential pathogenic variants with moderately reduced activity and trafficking efficiency; red, potential pathogenic variants with severely reduced activity and trafficking efficiency; magenta, the H84L variant; and grey, the C113S variant (a known pathogenic variant and positive control). The right panels show the zoomed-in views of key regions, including the center of the dimerization interface (upper panel) and the region surrounding H84 (lower panel).

**Figure 6.**
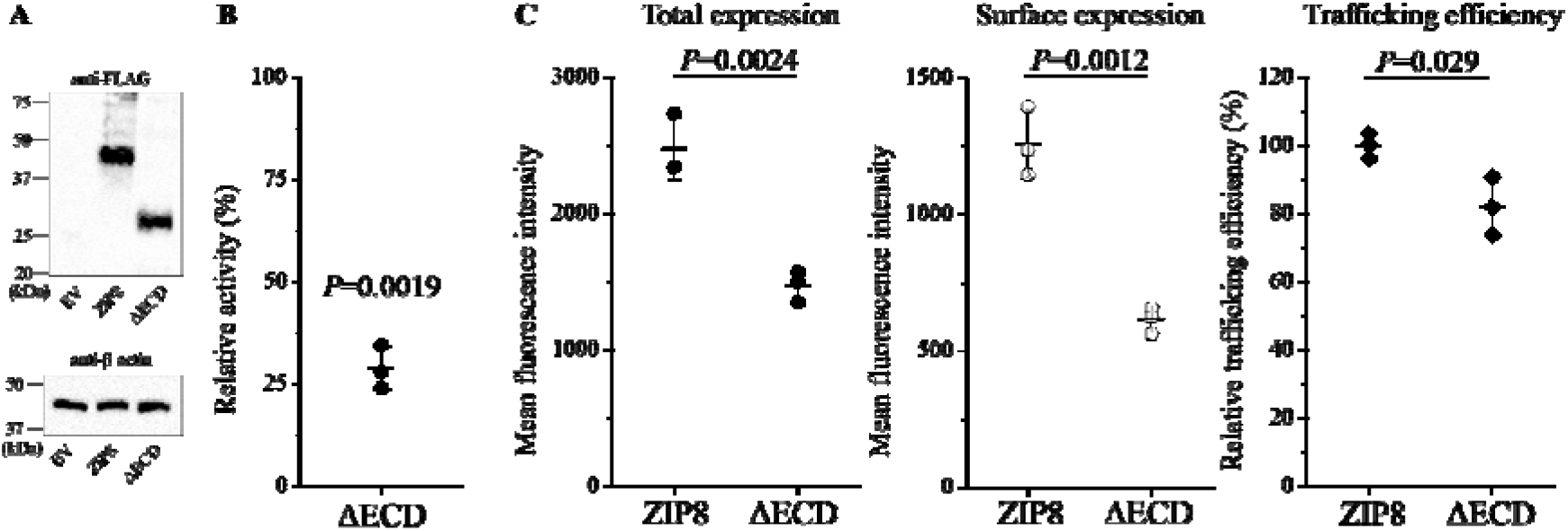
Functional characterization of the ΔECD variant of ZIP8 in comparison with wild-type ZIP8. (**A**) Western blot. N-terminal FLAG-tagged ZIP8 and its ΔECD variant were expressed in HEK293T cells and detected using the M2 anti-FLAG antibody in Western blot. β-actin was used as loading control. (**B**) Transport activity. The data shown are the mean activities, expressed as percentages of the activity of wild-type ZIP8, from three independent experiments. In each experiment, three biological replicates were tested. Two tailed one-sample t test was conducted to examine null hypothesis. (**C**) Total and cell surface expression quantified by flow cytometry. The data shown are from one of two independent experiments. In one experiment, three biological replicates were included for each condition. Two tailed t test was conducted to examine null hypothesis. In (B) and (C), the horizontal bars and error bars indicate the means and S.D.s, respectively.

To further address the function of ZIP8-ECD, we generated an N-terminal FLAG-tagged ZIP8 construct lacking the ECD (Δ23-116) and compared its transport activity, total expression, and cell surface expression with those of wild-type ZIP8. As shown in **Figure 6**, the ΔECD construct exhibited approximately 30% of the transport activity of full-length ZIP8. However, this drastic activity reduction can be largely attributed to the significantly reduced total expression and lower cell surface expression. A modest decrease in trafficking efficiency was also observed. These results suggest a potential role for the ZIP8-ECD in maintaining protein expression and promoting proper cellular trafficking. In contrast, ZIP4-ECD is crucial for optimal zinc transport (42). This function is primarily attributed to the N-terminal helix-rich domain, whereas the C-terminal PAL motif-containing domain, which is homologous to ZIP8-ECD, contributes only marginally to overall transport activity (36). Another notable difference between ZIP4 and ZIP8 is that deletion of the ZIP4-ECD increases total and cell surface expression, whereas ZIP8-ECD deletion decreases both total and cell surface expression. Together, these observations indicate that the ECDs of LIV-1 subfamily members can have distinct roles and varying degrees of importance in transporter activity.

## Conclusion

In this study, we applied an optimized LA-ICP-MS–based workflow that enables functional screening of a large number of variants with improved throughput. By integrating array-based sample preparation with 96-well plate cell assays, this method provides a robust and scalable platform for directly quantifying intracellular metal accumulation. We believe that this method can be readily extended to other members of the ZIP family, as well as to metal importers from distinct transporter families that require rapid quantitative measurement of metal uptake.

Using this method, we identified more than ten previously uncharacterized variants with significantly reduced transport activity, representing potential pathogenic mutations. These findings help bridge the gap between the growing number of reported genetic variants and the limited availability of experimental functional annotations. Our results also provide an opportunity to evaluate the performance of computational prediction tools. While AM shows reasonable predictive power and outperforms CADD in this dataset, the observed error rate of approximately 20% highlights the continued need for experimental validation. These data further emphasize the value of generating high-quality functional datasets to improve future prediction models.

Finally, the functional screening results offer new insights into the structure-function relationship of the ZIP8-ECD. The data suggest that impaired trafficking, likely arising from defects in protein folding and stability, accounts for the reduced activity observed in most functionally compromised variants. In addition, our analysis highlights the importance of the ZIP8-ECD in maintaining proper protein expression and cell surface localization, and suggests a potential role for ECD-mediated dimerization in transporter function. Together, these findings provide a more comprehensive understanding of how genetic variation impacts ZIP8 function and establish a generalizable framework for studying metal transporter variants.

## Experimental procedures

### Gene, plasmids, and reagents

As described previously (14,15), the complementary DNA of human ZIP8 (GenBank access number: BC012125) from Mammalian Gene Collection were purchased from GE Healthcare. The ZIP8 construct consists of the N-terminal signal peptide of ZIP4 (amino acid residues 1-22) followed by a GSGS linker and a FLAG tag, and the ZIP8 coding sequence (residue 23-460). Other reagents were purchased from Sigma-Aldrich or Fisher Scientific. Site-directed mutagenesis of ZIP8 was conducted using QuikChange mutagenesis kit (Agilent, Cat# 600250). All mutations were verified by DNA sequencing. The primers used in this study are listed in **Table S1**.

### Cell culture, transfection, and Western blot

Human embryonic kidney cells (HEK293T, ATCC, Cat# CRL-3216) were cultured in Dulbecco’s Modified Eagle Medium (DMEM, Thermo Fisher Scientific, Invitrogen, Cat# 11965092) supplemented with 10% (v/v) fetal bovine serum (FBS, Thermo Fisher Scientific, Invitrogen, Cat# 10082147). Cells were passaged at 70-90% confluency (every 2-3 days) and seeded in 96-well plates coated with poly-D-lysine at 10,000 cells/cm^2^. After growing to 70% confluency (24 hours), cells were transfected with plasmid DNA and lipofectamine 2000 (Thermo Fisher Scientific, Invitrogen, Cat# 11668019) following manufacturer’s instructions. For 96-well plates, 200 ng plasmid per well and 500 ng lipofectamine 2000 were used in each well.

For Western blot, samples were mixed with the SDS sample loading buffer and incubated at 37 °C for 20 min before loading on SDS-PAGE gel. The proteins separated by SDS-PAGE were transferred to polyvinylidene fluoride membranes (Millipore, Cat# PVH00010). After blocking with 5% (w/v) nonfat dry milk, the membranes were incubated with mouse M2 anti-FLAG antibody (Sigma, Cat# F3165) or rabbit anti-β-actin (Cell Signaling, Cat# 4970 S) at 4 °C overnight, which were detected with horseradish peroxidase-conjugated horse anti-mouse immunoglobulin-G antibody at 1:5000 dilution (Cell Signaling Technology, Cat# 7076), or goat anti-rabbit immunoglobulin-G antibody at 1:3000 dilution (Cell Signaling Technology, Cat# 7074), respectively, using the chemiluminescence reagent (VWR, Cat#RPN2232). The images of the blots were taken using a Bio-Rad ChemiDoc Imaging System.

### Cadmium uptake assay

Twenty hours post transfection, cells were washed with pre-warmed wash buffer (10 mM HEPES, 142 mM NaCl, 5 mM KCl, 10 mM glucose, pH 7.30). Wash buffer was removed and cadmium uptake solution (wash buffer supplemented with 2 mM CaCl_2_, 1 mM MgCl_2_, and 10 µM CdCl_2_) was added. Plates were placed in a 37 °C incubator for 1 h. After incubation, cells were put on ice, and an equal amount of ice-cold stop buffer (wash buffer plus 1 mM EDTA) was added, and then removed after a low speed centrifugation. Cells were washed three times with ice-cold wash buffer prior to sample array preparation.

### Sample array preparation for LA-ICP-MS

Glass slides were washed in soap and rinsed 5 times with Milli-Q water followed by drying overnight at 45 °C. The dried slides were coated with 5% dimethyldichlorosilane in heptane (Supelco, Cat# 85126) for 30 minutes at room temperature. After air-drying for a few hours, the slides were rinsed thoroughly with Milli-Q water (five times) and dried overnight prior to use. Cells were scraped and resuspended in 30 μl of Milli-Q water per well, and 7.5 μl of the cell suspension was applied onto the glass slides as separate droplets. The slides were then dried overnight in an incubator. When performed carefully, more than 50 samples can be accommodated on a single glass slide.

### LA-ICP-TOF-MS

As described previously (14), up to three sample arrays were loaded into a Bioimage 266 nm LA system (Elemental Scientific Lasers) which is equipped with an ultrafast low dispersion TwoVol3 ablation chamber and a dual concentric injector (DCI3) and is coupled to an icpTOF S2 (TOFWERK AG) ICP-TOF-MS. Using the optimized laser parameters (80% laser power, 9.5–10.5 J/cm^2^ laser fluence, 0.1–0.15 mJ sample energy) with a 40 μm spot size (circular) and repetition rate of 50 Hz, ablation of the glass slide was analyzed using ^27^Al and ^140^Ce elemental maps to ensure minimal glass ablation but complete sample ablation. Each sample spot was analyzed by ablating five parallel lines with 120 μm spacing between two adjacent lines. For each ablation line, the ^31^P counts were used to identify and exclude the data from the ablation points where the cell lysate content is very low. Then, the counts per minute of ^31^P and ^111^Cd for each sample spot were integrated in the regions drawn manually for each spot, respectively, and used to calculate the ^111^Cd/^31^P count ratio.

### Flow cytometry

Total and cell surface expression levels of any variant were determined using cells from the same experiment. To measure cell surface expression of FLAG-tagged ZIP8 variants, cells were trypsinized, resuspended, and fixed with 4% formaldehyde at room temperature for 15 min. After a brief wash, cells were incubated with anti-FLAG antibody (Sigma, Cat# F3165) at a 1:500 dilution in Dulbecco’s phosphate-buffered saline (DPBS) for 1 h on ice. To determine total expression levels, the same procedure was followed except that cells were fixed with 4% formaldehyde containing 0.1% Triton X-100 to allow permeabilization. After the unbound antibodies were removed by washing with DPBS, the cells were incubated with an Alexa Fluor 568-conjugated goat anti-mouse antibody (Invitrogen, CAT#A-11004) at 1 : 1000 diluted in DPBS for 2 h. After washing three times with a DPBS without Ca^2+^ and Mg^2+^, cells were re-suspended with DPBS for flow cytometry analysis using a ThermoFisher Attune Cytpix flow cytometer. at least 10,000 cells were assessed for each condition. Single cells were selected and analyzed to detect the fluorescence of Alexa Fluor 568. The cellular debris was excluded based on FSC-A vs. SSC-A panel and single cells were selected through FSC-A vs FSC-H gating (**Figure S4A**). The mean fluorescence intensity was calculated for each sample and representative results are shown in **Figure S4B**. Data processing was performed in FlowJo 10.8.1 and GraphPad Prism 10.

## Quantification and statistical analysis

We assumed a normal distribution of the samples and significant difference were examined using two-tailed, paired, or one-sample Student’s *t*-test as indicated. A *P*< 0.05 was considered statistically significant. Data were presented as mean ± standard deviation.

## Data availability statement

Further information and requests for resources and reagents, including the plasmids generated in this study, should be directed to Dr. Jian Hu. Any additional information required to reanalyze the data reported in this paper is available from Dr. Jian Hu upon request.

## Acknowledgement

The data presented herein were obtained using instrumentation in the MSU Flow Cytometry Core Facility. The facility is funded in part through the financial support of Michigan State University’s Office of Research & Innovation and Colleges of Osteopathic Medicine, Human Medicine, Veterinary Medicine, Natural Sciences, and Engineering. The Attune CytPix is supported by the Equipment Grants Program, award #2022-70410-38419, from the U.S. Department of Agriculture, National Institute of Food and Agriculture. We thank Quantitative Bio Element Analysis and Mapping (QBEAM) center at Michigan State University for the assistance of using mass spectrometry. This work was supported by NIH grants GM140931 and TR005599 (to J.H.) and P41-GM135018, GM115848, and GM038784 (to T.O.).

## Author contributions

Investigation: M.K., T.W., A.S., K.M., and H.Z; Writing: M.K., T.W., A.S., K.M., H.Z., T.O., and J.H.; Conceptualization: T.V.O. and J.H.; Supervision: T.V.O. and J.H.

## Conflict of Interest Statement

The authors declare no competing interests.

## Notes

### Competing Interest Statement

The authors have declared no competing interest.

