## Supplementary Information for "Functional screening of ZIP8 naturally occurring variants identifies pathogenic mutations and trafficking defects"

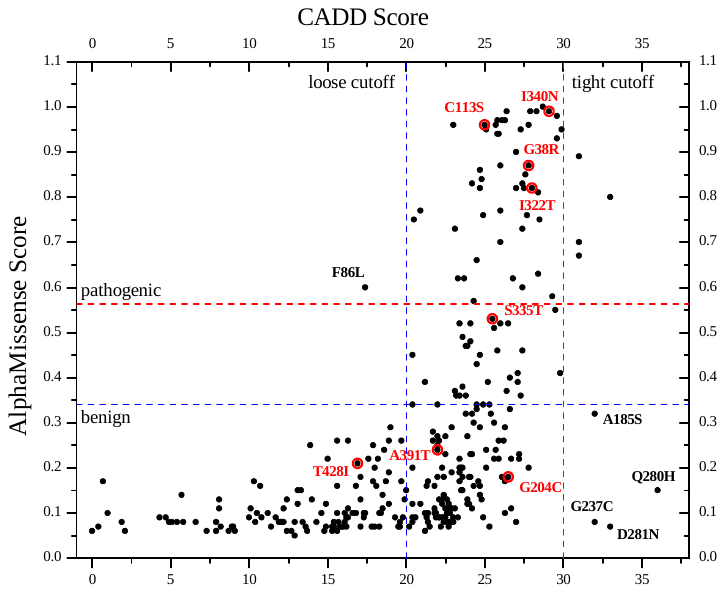

**Figure S1.** Pathogenicity predictions of naturally occurring non-somatic variants for human ZIP8. For each selected variant, the AlphaMissense (AM) pathogenicity score is plotted against the CADD score. The red circles indicate confirmed pathogenic variants. The dashed lines indicate the cutoffs: for CADD, the loose cutoff is set at 20, and the tight cutoff is at 30; for AM, the cutoffs for pathogenic and benign are 0.564 and 0.34, respectively. The confirmed pathogenic variants and those with drastically conflicting prediction scores are labeled.

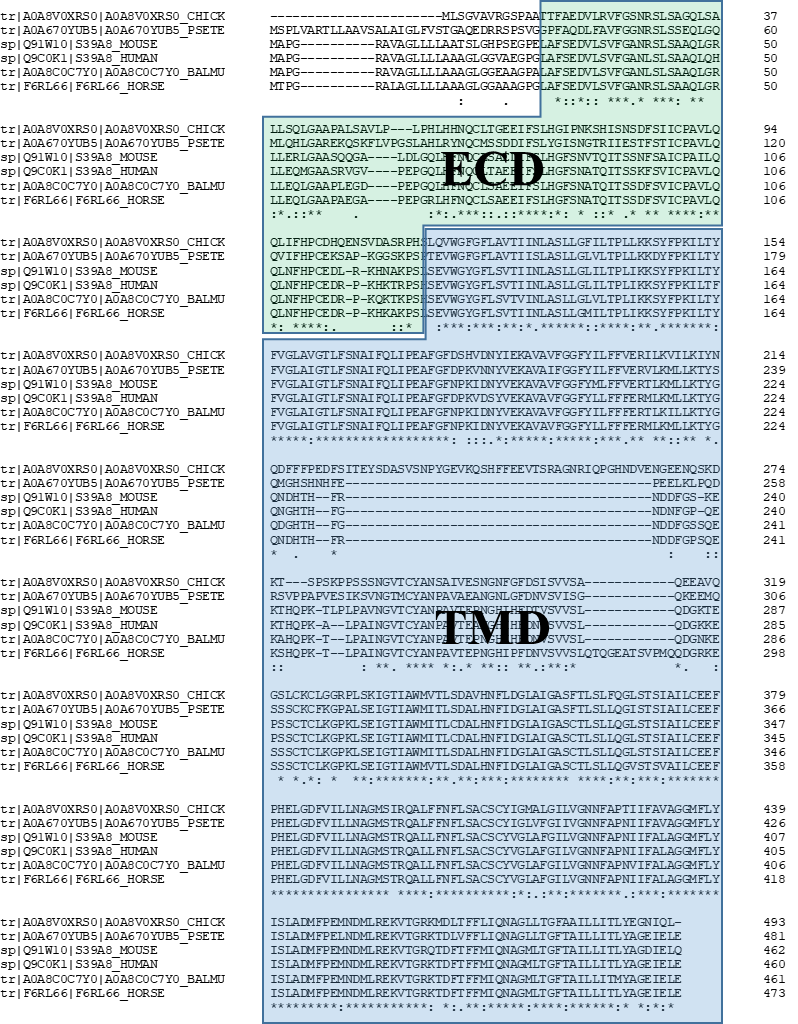

**Figure S2.** Multiple sequence alignment of human ZIP8 and its homologs. The ECD and TMD regions are highlighted in the green and blue frames, respectively, and the ECD is less conserved than the TMD.

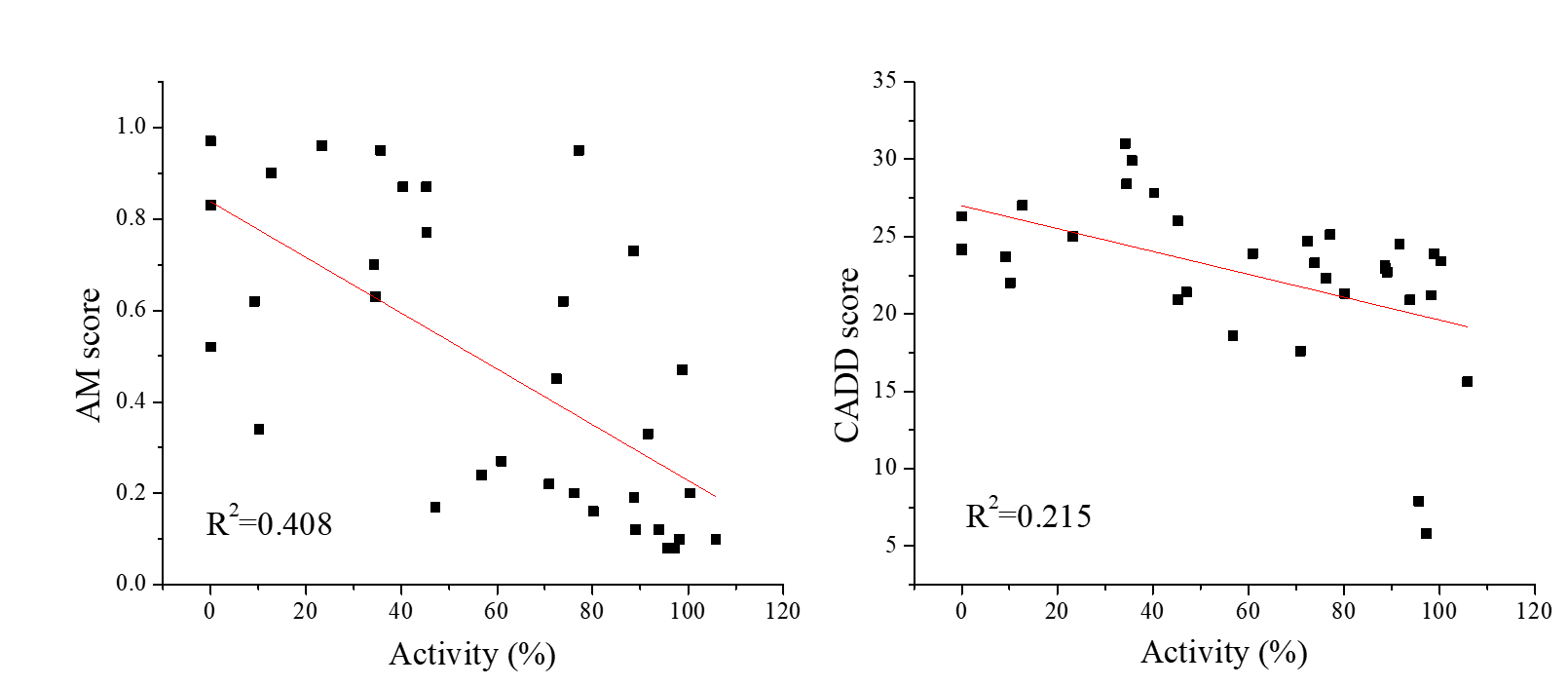

**Figure S3.** Correlation of variant activity with AlphaMissense pathogenicity score (*left*) or with CADD score (*right*). The coefficients of determination (R^2^) of the linear correlations are indicated in each plot.

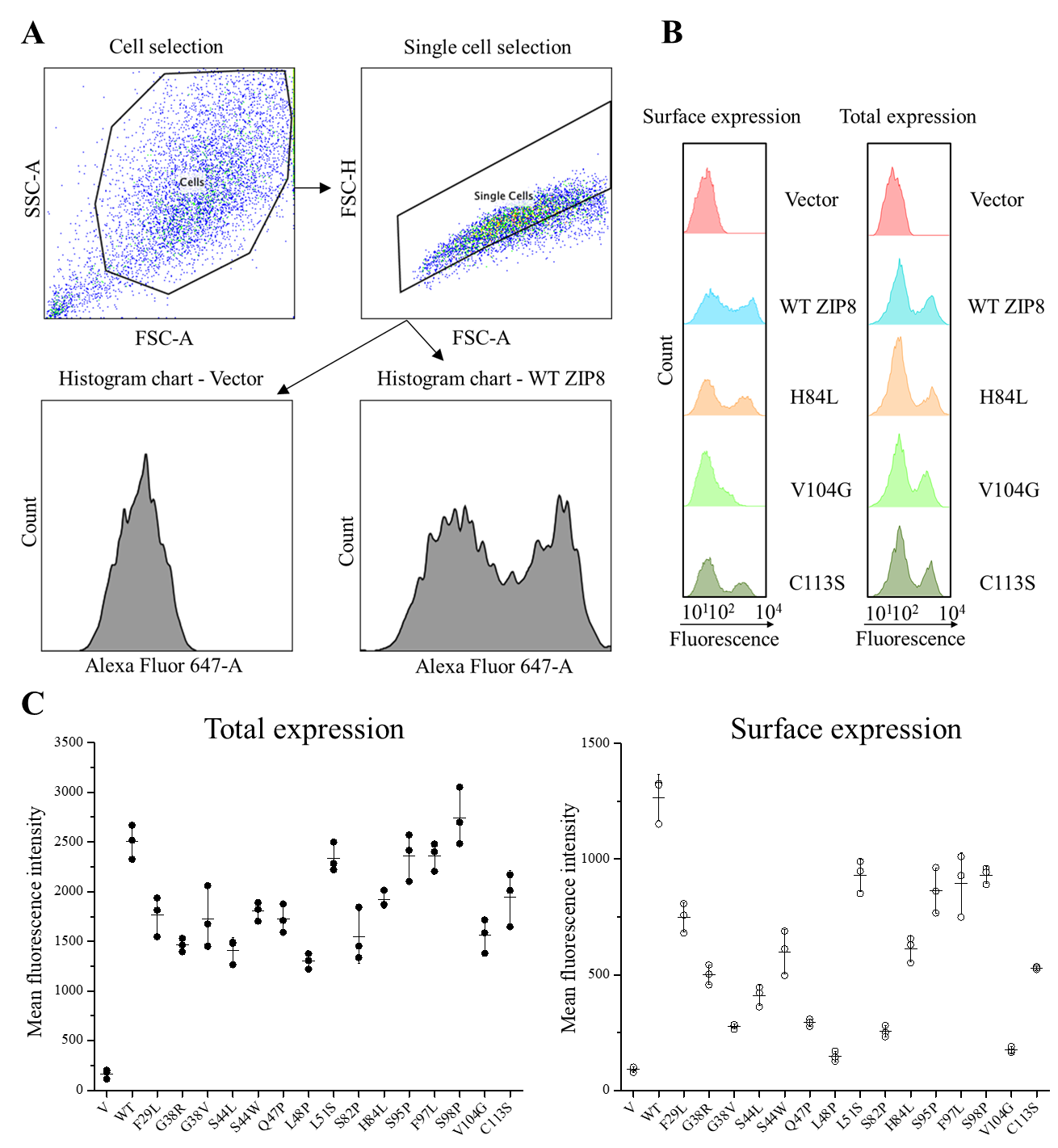

**Figure S4.** Processing and analysis of flow cytometry data. (**A**) Flow chart of flow cytometry data processing. (**B**) Histogram plots of flow cytometry data for vector, ZIP8, and representative variants. (**C**) Mean fluorescence intensities for the total expression (*left*) and cell surface expression (*right*) of vector, wild-type ZIP8, and the potential pathogenic variants studied in this work. The data shown is from one of two independent experiments and three biological replicates were conducted for each variant in one experiment. The horizontal bars and error bars indicate the mean and S.D., respectively.

**Table S1.** Primers used in this study.

| F29L-F: F29L-R: | 5’ TCC GAG GGG CCA GGG CTA GCC CTG AGC GAG GAT GTG CTG AGC GTG 3’  5’ CAC GCT CAG CAC ATC CTC GCT CAG GGC TAG CCC TGG CCC CTC GGA 3’ |
| --- | --- |
| D32E-F: D32E-R: | 5’ CCA GGG CTA GCC TTC AGC GAG GAG GTG CTG AGC GTG TTC GGC GCG 3’  5’ CGC GCC GAA CAC GCT CAG CAC CTC CTC GCT GAA GGC TAG CCC TGG 3’ |
| V33M-F: V33M-R: | 5’ GGG CTA GCC TTC AGC GAG GAT ATG CTG AGC GTG TTC GGC GCG AAT 3’  5’ ATT CGC GCC GAA CAC GCT CAG CAT ATC CTC GCT GAA GGC TAG CCC 3’ |
| V36G-F: V36G-R: | 5’ TTC AGC GAG GAT GTG CTG AGC GGC TTC GGC GCG AAT CTG AGC CTG 3’  5’ CAG GCT CAG ATT CGC GCC GAA GCC GCT CAG CAC ATC CTC GCT GAA 3’ |
| F37L-F: F37L-R: | 5’ AGC GAG GAT GTG CTG AGC GTG CTG GGC GCG AAT CTG AGC CTG TCG 3’  5’ CGA CAG GCT CAG ATT CGC GCC CAG CAC GCT CAG CAC ATC CTC GCT 3’ |
| G38R-F: G38R-R: | 5’ GAG GAT GTG CTG AGC GTG TTC CGG GCG AAT CTG AGC CTG TCG GCG 3’  5’ CGC CGA CAG GCT CAG ATT CGC CCG GAA CAC GCT CAG CAC ATC CTC 3’ |
| G38V-F: G38V-R: | 5’ GAG GAT GTG CTG AGC GTG TTC GTT GCG AAT CTG AGC CTG TCG GCG 3’  5’ CGC CGA CAG GCT CAG ATT CGC AAC GAA CAC GCT CAG CAC ATC CTC 3’ |
| S42R-F: S42R-R: | 5’ AGC GTG TTC GGC GCG AAT CTG CGG CTG TCG GCG GCG CAG CTC CAG 3’  5’ CTG GAG CTG CGC CGC CGA CAG CCG CAG ATT CGC GCC GAA CAC GCT 3’ |
| L43V-F: L43V-R: | 5’ GTG TTC GGC GCG AAT CTG AGC GTG TCG GCG GCG CAG CTC CAG CAC 3’  5’ GTG CTG GAG CTG CGC CGC CGA CAC GCT CAG ATT CGC GCC GAA CAC 3’ |
| S44L-F: S44L-R: | 5’ TTC GGC GCG AAT CTG AGC CTG CTG GCG GCG CAG CTC CAG CAC TTG 3’  5’ CAA GTG CTG GAG CTG CGC CGC CAG CAG GCT CAG ATT CGC GCC GAA 3’ |
| S44W-F: S44W-R: | 5’ TTC GGC GCG AAT CTG AGC CTG TGG GCG GCG CAG CTC CAG CAC TTG 3’  5’ CAA GTG CTG GAG CTG CGC CGC CCA CAG GCT CAG ATT CGC GCC GAA 3’ |
| A45S-F: A45S-R: | 5’ GGC GCG AAT CTG AGC CTG TCG AGC GCG CAG CTC CAG CAC TTG CTG 3’  5’ CAG CAA GTG CTG GAG CTG CGC GCT CGA CAG GCT CAG ATT CGC GCC 3’ |
| Q47P-F: Q47P-R: | 5’ AAT CTG AGC CTG TCG GCG GCG CCC CTC CAG CAC TTG CTG GAG CAG 3’  5’ CTG CTC CAG CAA GTG CTG GAG GGG CGC CGC CGA CAG GCT CAG ATT 3’ |
| L48P-F: L48P-R: | 5’ CTG AGC CTG TCG GCG GCG CAG CCC CAG CAC TTG CTG GAG CAG ATG 3’  5’ CAT CTG CTC CAG CAA GTG CTG GGG CTG CGC CGC CGA CAG GCT CAG 3’ |
| L51S-F  L51S-R: | 5’ TCG GCG GCG CAG CTC CAG CAC AGC CTG GAG CAG ATG GGA GCC GCC 3’  5’ GGC GGC TCC CAT CTG CTC CAG GCT GTG CTG GAG CTG CGC CGC CGA 3’ |
| M55R-F:  M55R-R: | 5’ CTC CAG CAC TTG CTG GAG CAG CGG GGA GCC GCC TCC CGC GTG GGC 3’  5’ GCC CAC GCG GGA GGC GGC TCC CCG CTG CTC CAG CAA GTG CTG GAG 3’ |
| M55T-F:  M55T-R: | 5’ CTC CAG CAC TTG CTG GAG CAG ACC GGA GCC GCC TCC CGC GTG GGC 3’  5’ GCC CAC GCG GGA GGC GGC TCC GGT CTG CTC CAG CAA GTG CTG GAG 3’ |
| A58D-F:  A58D-R: | 5’ TTG CTG GAG CAG ATG GGA GCC GAC TCC CGC GTG GGC GTC CCG GAG 3’  5’ CTC CGG GAC GCC CAC GCG GGA GTC GGC TCC CAT CTG CTC CAG CAA 3’ |
| P64S-F:  P64S-R: | 5’ GCC GCC TCC CGC GTG GGC GTC AGC GAG CCT GGC CAG CTG CAC TTC 3’  5’ GAA GTG CAG CTG GCC AGG CTC GCT GAC GCC CAC GCG GGA GGC GGC 3’ |
| E65D-F:  E65D-R: | 5’ GCC TCC CGC GTG GGC GTC CCG GAC CCT GGC CAG CTG CAC TTC AAC 3’  5’ GTT GAA GTG CAG CTG GCC AGG GTC CGG GAC GCC CAC GCG GGA GGC 3’ |
| L69R-F: L69R-R: | 5’ GGC GTC CCG GAG CCT GGC CAG CGT CAC TTC AAC CAG TGT TTA ACT 3’  5’ AGT TAA ACA CTG GTT GAA GTG ACG CTG GCC AGG CTC CGG GAC GCC 3’ |
| N72S-F:  N72S-R: | 5’ GAG CCT GGC CAG CTG CAC TTC AGC CAG TGT TTA ACT GCT GAA GAG 3’  5’ CTC TTC AGC AGT TAA ACA CTG GCT GAA GTG CAG CTG GCC AGG CTC 3’ |
| S82F-F:  S82F-R: | 5’ TTA ACT GCT GAA GAG ATC TTT TTC CTT CAT GGC TTT TCA AAT GCT 3’  5’ AGC ATT TGA AAA GCC ATG AAG GAA AAA GAT CTC TTC AGC AGT TAA 3’ |
| S82P-F:  S82P-R: | 5’ TTA ACT GCT GAA GAG ATC TTT CCC CTT CAT GGC TTT TCA AAT GCT 3’  5’ AGC ATT TGA AAA GCC ATG AAG GGG AAA GAT CTC TTC AGC AGT TAA 3’ |
| H84L-F:  H84L-R: | 5’ GCT GAA GAG ATC TTT TCC CTT CTG GGC TTT TCA AAT GCT ACC CAA 3’  5’ TTG GGT AGC ATT TGA AAA GCC CAG AAG GGA AAA GAT CTC TTC AGC 3’ |
| Q91L-F: Q91L-R: | 5’ CAT GGC TTT TCA AAT GCT ACC CTG ATA ACC AGC TCC AAA TTC TCT 3’  5’ AGA GAA TTT GGA GCT GGT TAT CAG GGT AGC ATT TGA AAA GCC ATG 3’ |
| S95P-F: S95P-R: | 5’ AAT GCT ACC CAA ATA ACC AGC CCC AAA TTC TCT GTC ATC TGT CCA 3’  5’ TGG ACA GAT GAC AGA GAA TTT GGG GCT GGT TAT TTG GGT AGC ATT 3’ |
| F97L-F:  F97L-R: | 5’ ACC CAA ATA ACC AGC TCC AAA CTG TCT GTC ATC TGT CCA GCA GTC 3’  5’ GAC TGC TGG ACA GAT GAC AGA CAG TTT GGA GCT GGT TAT TTG GGT 3’ |
| S98P-F: S98P-R: | 5’ CAA ATA ACC AGC TCC AAA TTC CCC GTC ATC TGT CCA GCA GTC TTA 3’  5’ TAA GAC TGC TGG ACA GAT GAC GGG GAA TTT GGA GCT GGT TAT TTG 3’ |
| P102A-F: P102A-R: | 5’ TCC AAA TTC TCT GTC ATC TGT GCG GCA GTC TTA CAG CAA TTG AAC 3’  5’ GTT CAA TTG CTG TAA GAC TGC CGC ACA GAT GAC AGA GAA TTT GGA 3’ |
| V104G-F:  V104G-R: | 5’ TTC TCT GTC ATC TGT CCA GCA GGC TTA CAG CAA TTG AAC TTT CAC 3’  5’ GTG AAA GTT CAA TTG CTG TAA GCC TGC TGG ACA GAT GAC AGA GAA 3’ |
| F110I-F: F110I-R: | 5’ GCA GTC TTA CAG CAA TTG AAC ATC CAC CCA TGT GAG GAT CGG CCC 3’  5’ GGG CCG ATC CTC ACA TGG GTG GAT GTT CAA TTG CTG TAA GAC TGC 3’ |
| C113S-F C113S-R: | 5’ CAG CAA TTG AAC TTT CAC CCA AGC GAG GAT CGG CCC AAG CAC AAA 3’  5’ TTT GTG CTT GGG CCG ATC CTC GCT TGG GTG AAA GTT CAA TTG CTG 3’ |
| ZIP8-F: | 5’ TAT ATA AGC AGA GCT 3’ (for sequencing) |
| ZIP8-ECD-F: | 5’ GATTACAAGGACGACGATGACAAGGGATCCCCCAAGCACAAAACAAGACCA AGTCATTCA 3’ |
| ZIP8-ECD-R: | 5’ TGAATGACTTGGTCTTGTTTTGTGCTTGGGGGATCCCTTGTCATCGTCGTCC  TTGTAATC 3’ |
